# CRISPR/Cas9 generated *MeSWEET10a* mutants show reduced susceptibility to cassava bacterial blight and produce viable seed

**DOI:** 10.1101/2023.06.21.545993

**Authors:** Kiona Elliott, Kira M. Veley, Greg Jensen, Kerrigan B. Gilbert, Joanna Norton, Lukas Kambic, Marisa Yoder, Alex Weil, Sharon Motomura-Wages, Rebecca S. Bart

**Affiliations:** Donald Danforth Plant Science Center; Saint Louis, MO 63132, USA; Division of Biological and Biomedical Sciences, Washington University in Saint Louis, St. Louis, MO, 63110, USA; College of Tropical Agriculture & Human Resources, University of Hawaii at Manoa, Hilo, HI 96720, USA

**Author notes:** Corresponding author. Email: RSB.

**Keywords:** Cassava bacterial blight, *Xanthomonas*, SWEET gene, TAL effector, CRISPR/Cas9, Genome editing

## Abstract

Bacteria from the genus *Xanthomonas* are prolific phytopathogens that elicit disease in over 400 plant species. Xanthomonads carry a repertoire of specialized proteins called transcription activator-like (TAL) effectors that promote disease and pathogen virulence by inducing expression of host susceptibility (S) genes. *Xanthomonas phaseoli* pv. *manihotis* (Xpm) causes bacterial blight on the staple food crop, cassava. The Xpm effector, TAL20, induces ectopic expression of the S gene, *MeSWEET10a*, a sugar transporter that contributes to cassava bacterial blight susceptibility. We used CRISPR/Cas9 to generate multiple cassava lines with edits to the *MeSWEET10a* TAL20 effector binding site and/or coding sequence. In several of the regenerated lines, *MeSWEET10a* expression was no longer induced by *Xpm* and in these cases, we observed reduced cassava bacterial blight disease symptoms post Xpm infection. *MeSWEET10a* is expressed in cassava flowers. Therefore, we investigated flower development and reproductive function in an *MeSWEET10a* mutant line. We found that the *MeSWEET10a* mutant produced phenotypically wildtype cassava flowers and viable F1 seed. Thus, blocking *MeSWEET10a* induction is a viable strategy for decreasing cassava susceptibility to CBB.

## Introduction

Cassava (*Manihot esculenta* Crantz) is a starchy root crop that serves as a carbohydrate source and food security crop for nearly 800 million people globally [(Alves, 2002) and (Morgan and Choct, 2016)], is tolerant to abiotic stressors and is often grown without costly inputs like fertilizer (EL-Sharkawy, 2003). This crop is especially important for smallholder farmers in Sub-Saharan Africa who grow cassava as a sustenance crop and sell it for revenue when yields allow [(Taylor et al., 2003) and (Hillocks et al., 2002)]. A leading biotic factor threatening cassava production is cassava bacterial blight (CBB) (Lozano, 1986). CBB disease symptoms include water-soaked leaf lesions, chlorosis, defoliation, and stem browning (Lozano, 1986). CBB is present in all cassava growing regions and can result in total crop loss including the stem used to plant a subsequent crop through clonal propagation [(Lozano et al., 1980) and (López and Bernal, 2012)].

The causal agent of CBB is a gram-negative phytopathogen in the genus *Xanthomonas*. Xanthomonads elicit disease in over 400 plant species including economically important crops such as rice, cotton, sorghum, and citrus [(Leyns et al., 1984), (Jacques et al., 2016), and (Mhedbi-Hajri et al., 2013)]. The *Xanthomonas* specific to cassava was recently reclassified as *X. phaseoli* pv. *manihotis* (Xpm) (Constantin et al., 2016) and was formerly known as *X. axonopodis* pv. *manihotis* (Xam). Xpm is dispersed from plant to plant through rain, wind, or by the propagation of already infected stem cuttings. From the leaf surface, Xpm can enter the leaf through open stomata or wounds (Kandel et al., 2017). *In planta, Xanthomonas* colonizes the surface of mesophyll cells and some Xanthomonads, including Xpm, can systemically spread throughout the plant vasculature [(An et al., 2019) and (Ryan et al., 2011)].

Xpm induces effector triggered susceptibility (ETS) using an arsenal of effector proteins released into the plant through a needle-like projection that penetrates the host cell wall called the type III secretion system (T3SS) [(Bart et al., 2012) and (Abrusci et al., 2014)]. T3SS effectors manipulate the host to help the pathogen overcome plant defenses and promote disease (Hogenhout et al., 2009). Bacteria in the *Xanthomonas* and *Ralstonia* genera have specialized transcription activator-like (TAL) effectors that induce expression of host susceptibility (S) genes to enhance pathogenesis [(van Schie and Takken, 2014) and (Eckardt, 2002)]. TAL effectors structurally resemble eukaryotic transcription factors with an activation domain, nuclear localization signal, and a DNA binding domain consisting of tandem amino acid repeats (Schornack et al., 2013). The DNA binding domain directs the TAL effector to predictable DNA sequences, called effector binding elements (EBEs) [(Cernadas et al., 2014), (Moscou and Bogdanove, 2009), and (Boch and Bonas, 2010)]. In many cases, TAL effectors binding causes upregulation of downstream susceptibility genes.

*Xpm* strains typically carry between one to five TAL effectors and the model Xpm strain used in this study, Xpm668 (formerly known as Xam668), has five TAL effectors: TAL13, TAL14, TAL15, TAL20, and TAL22 (Cohn and Bart et al., 2014). Xpm TAL20 mutants (Xpm668ΔTal20) show reduced virulence with the most obvious phenotype being a reduction in the water-soaked lesions that are typical of this disease (Cohn and Bart et al., 2014). A member of the SWEET (Sugars Will Eventually be Exported Transporter) gene family, *MeSWEET10a* (Gene ID: Manes.06G123400), was identified as the S gene target for TAL20. TAL20 binding to the *MeSWEET10a* EBE induces ectopic gene expression in the leaf and results in sugar transport into the apoplast where Xpm proliferates (Cohn and Bart et al., 2014). SWEET genes have been studied as pathogen virulence factors in other pathosystems [(Chen, 2014) and (Chen et al., 2010)]. Furthermore, SWEET genes are established TAL effector targets in several plant species including rice, pepper, and cotton [(Antony et al., 2010), (Hu et al., 2014), (Phillips et al., 2017), and (Cox et al., 2017)] and in several cases, preventing TAL effector binding reduced plant susceptibility to disease [(Gupta et al., 2021), (Veley et al., 2023) and (Oliva et al., 2019)]. Additional classes of susceptibility genes have been described from various systems, for example, *Cs LATERAL ORGAN BOUNDARIES 1 (LOB1)*, which is a TAL-induced target of citrus canker in sweet orange (Huang et al., 2022).

In this study, we used a dual gRNA CRISPR/Cas9 strategy to generate *MeSWEET10a* mutant lines with edits to the TAL20 EBE and/or gene coding sequence. We characterized the disease phenotypes of Xpm infected plants and demonstrated that *MeSWEET10a* mutants exhibit reduced cassava bacterial blight symptoms. Additionally, while *MeSWEET10a* is not normally expressed in cassava leaves, prior work showed there is endogenous expression in flower tissue [(Veley et al., 2021) and (Perera et al., 2012)]. In rice, knocking out the SWEET gene, *OsSWEET15*, led to reduced rice fertility (Hu et al., 2023). Therefore, we investigated the impact of editing *MeSWEET10a* on cassava flower development and reproductive function. We found that *MeSWEET10a* mutant cassava plants developed flowers morphologically similar to wildtype plants based on macro imaging. When these flowers were used for crosses, they produced fruit, and viable F1 seed.

## Results

We hypothesized that editing *MeSWEET10a* would reduce cassava susceptibility to *Xpm*. To test this hypothesis, we designed a single CRISPR/Cas9 construct (construct 108) with two guide RNAs (gRNAs), gRNA1 and gRNA2, which target the TAL20 EBE site and the translation start site (Start Codon: ATG), respectively. Additionally, while previous reports demonstrate that there is low efficiency of HDR in plants [(Puchta, 2005) and (Britt and May, 2003)], we optimistically included a repair template with homology arms that flank the EBE to allow for potential CRISPR-mediated homology-directed repair (HDR). The repair template was designed to replace the EBE with a sequence that TAL20 would not bind while maintaining the annotated TATA box (**Figure 1A**). Agrobacterium-mediated transformation was carried out in friable embryogenic callus (FEC) from the farmer-preferred cultivar of cassava, TME419, also referred to as WT419 (Chauhan et al., 2015). In total, thirty transgenic lines were recovered. The *MeSWEET10a* region of interest was amplified from each recovered transgenic line. Restriction digest was used to identify lines with potential EBE repair template integration and larger INDELs. If the EBE repair template was integrated, we expected it to abolish an HaeIII restriction enzyme site at the gRNA2 repair template site (**Supplemental Figure 1A**). Based on this analysis, transgenic lines with integration of the repair template were not recovered. However, several lines exhibited digest patterns different from WT419 (for example, lines #269 and #338, **Supplemental Figure 1B**). These lines were genotyped using Sanger sequencing along with line #2, which showed a wildtype-like digest pattern (**Supplemental Figure 2A**). The genotyping revealed that line #2 is a WT-like transgenic. Line #243 contains a five-base pair (bp) deletion on one allele and a 156 bp deletion on the other allele, both in the promoter region, which may affect TAL effector binding and/or expression. Line #323 is also biallelic, but the mutations are not predicted to affect TAL effector binding or expression of the gene. Line #269 has a large 122 bp deletion including the TATA Box, TAL20 EBE, and the *MeSWEET10a* start codon consistent with the observed smaller PCR product (**Figure 1D**). Line #338 has a 5 bp deletion upstream of the TATA box and TAL20 EBE and a 13 bp deletion after the start codon causing a frameshift and stop codon in exon 1. Lines #269 and #338 were confirmed as homozygous mutants using genomic DNA (gDNA) clone sequencing (**Figure 1C** and **Supplemental Figure 2B**).

**Figure 1:**
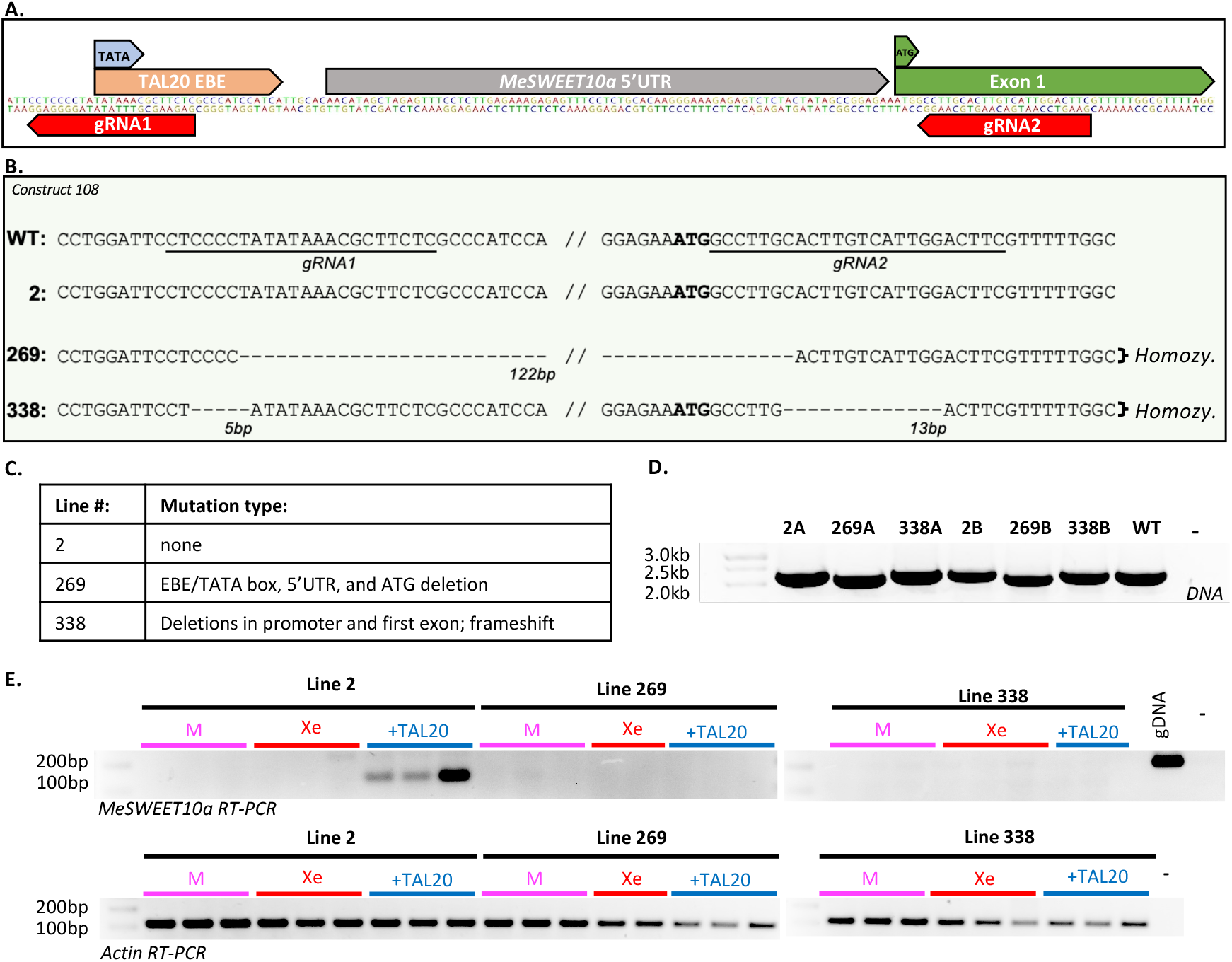
First generation *MeSWEET10a m*utant lines lack TAL20 mediated induction. **A)** Graphic depicting the *MeSWEET10a* region of interest including the TAL20 EBE (effector binding element), TATA box, translation start site (ATG), and gRNA target sites for construct 108. **B)** Genotyping of *MeSWEET10a* mutant lines recovered from construct 108 based on Sanger-sequencing. Text indicates sequences at the region of interest for wildtype plants and mutant lines #2, #269, and #338. Underlined text denotes the target site for each gRNA. Deletions are indicated by ‘-’ and the number of deleted base pairs (bp) is indicated below each deletion. Deletions in lines #269 and #338 are homozygous (homozyg.) **C)** Table with description of mutation and location type for each mutant line. **D)** PCR products generated by primers targeting the *MeSWEET10a* region in gDNA from WT and lines #2, #269, and #338. A and B denote different individuals from each line. **E)** RT-PCR of wildtype cassava and *MeSWEET10a* mutant lines infiltrated with mock, *X. euvesicatoria* (Xe) alone, and Xe+TAL20 treatments. Top gel shows results of RT-PCR with primers amplifying *MeSWEET10a* with an expected product size of 123 bp. Bottom gel shows results of RT-PCR with primers amplifying the housekeeping gene, *Actin*, as a control for sample loading with an expected product size of 125 bp. DNA from WT419 leaf tissue is included as a positive control and ‘-’ denotes a negative water control. M=Mock (magenta), Xe=(red), and +TAL20=Xe+TAL20 (blue).

**Figure 2:**
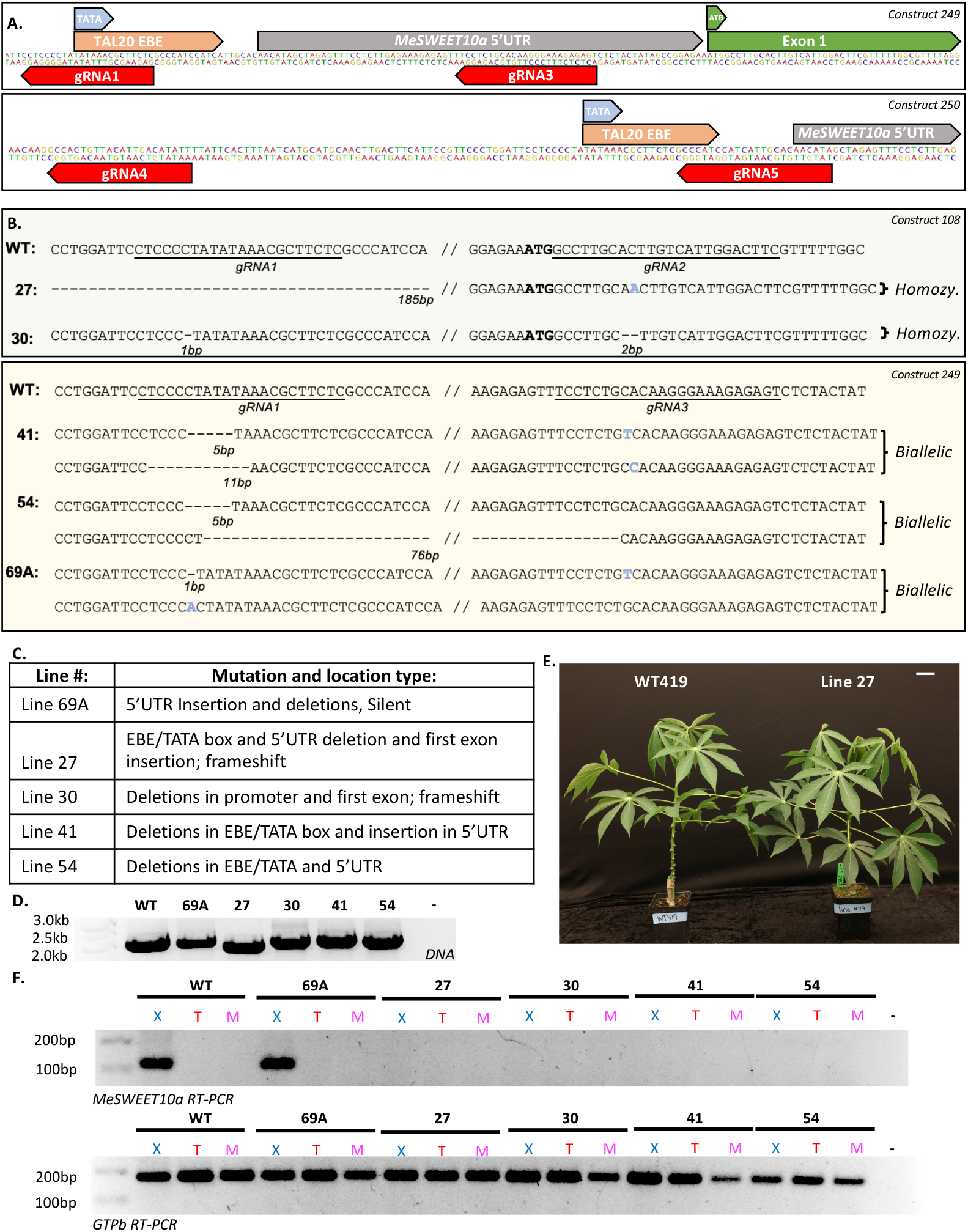
Additional *MeSWEET10a* mutant lines lack TAL20 mediated induction. A) Graphic depicting the *MeSWEET10a* region of interest, TAL20 EBE site, TATA box, translation start site (ATG), and gRNA target sites for constructs 108, 249, and 250. **B)** Genotyping of *MeSWEET10a* mutant lines recovered from construct 108 and 249 based on Sanger-sequencing. Text indicates sequences at the region of interest for wildtype plants and mutant lines #27, #30, #41, #54, and #69A. Underlined text denotes the gRNA target site. The italicized text below the underlined text describes gRNA number. The number of base pair (bp) deletions at each target site is depicted at the end of the sequence. Bases in blue indicate insertion events. Mutant line zygosity is indicated by ‘Homozy’ (homozygous) or ‘biallelic’ text to the right of the sequence. **C)** Table with description of mutation and location type for each mutant line. **D)** PCR products generated by primers targeting the *MeSWEET10a* region in gDNA from WT and edited lines. **E)** Representative image of wildtype cassava (left) and line #27 (right) plants grown from stake cuttings in greenhouse. Scale bar = 14cm. **F)** RT-PCR of wildtype cassava and *MeSWEET10a* mutant lines infected with Xpm, Xpm△TAL20, and mock treatments. The top gel shows results of RT-PCR with primers amplifying *MeSWEET10a* with an expected product size of 123 bp. The bottom gel shows results of RT-PCR with primers amplifying the housekeeping gene *GTPb* as a control for sample loading with an expected product size of 184 bp. ‘-’ denotes a negative water control. X= Xpm (blue), T=Xpm△TAL20 (red), and M=Mock (magenta).

We previously demonstrated that a TAL effector-less Xanthomonad, *Xanthomonas euvesicatoria* (Xe-non-pathogenic to cassava), can deliver TAL20 to cassava cells and induce *MeSWEET10a* expression (Cohn and Bart et al., 2014). We used this system to compare Xe, Xe +TAL20, or mock treatments for *MeSWEET10a* induction in lines #2, #269, and #338. At 48 hours post infection (HPI), samples were collected for RNA extraction and reverse transcription (RT)-PCR (**Figure 1E**). In control line #2 plants infected with Xe +TAL20, RT-PCR results show a 123 bp product indicating TAL20-mediated induction of *MeSWEET10a*. In contrast, no product was present for plants infected with Xe alone or mock treatments. Mutant lines #269 and #338 infected with Xe +TAL20 have no RT-PCR product indicating the mutations in each line are sufficient to prevent TAL20-mediated induction of *MeSWEET10a*.

Two additional CRISPR/Cas9 constructs were tested (**Figure 2A**). Construct 249 contains gRNA1 and gRNA3 which localize to the TAL20 EBE site and the *MeSWEET10a* 5’UTR (untranslated region) upstream of the start codon. Construct 250 contains gRNA4 which targets upstream of the TATA box and TAL20 EBE and gRNA5 which targets the TAL20 EBE downstream of the TATA box. Four rounds of cassava transformation with all three constructs were performed. In total, twenty-four transgenic lines were recovered with seven mature lines generated from construct 108, eight from construct 249, and nine from construct 250. Leaf tissue was sampled from each line at the plantlet stage in tissue culture and plants were genotyped by Sanger sequencing. Twenty-three out of twenty-four lines had edits within the *MeSWEET10a* promoter and/or coding sequence (**Supplemental Figure 3A**). One line was recovered containing only edits within the TAL20 binding site while maintaining an intact TATA box, however, this line died during the tissue culture process.

**Figure 3:**
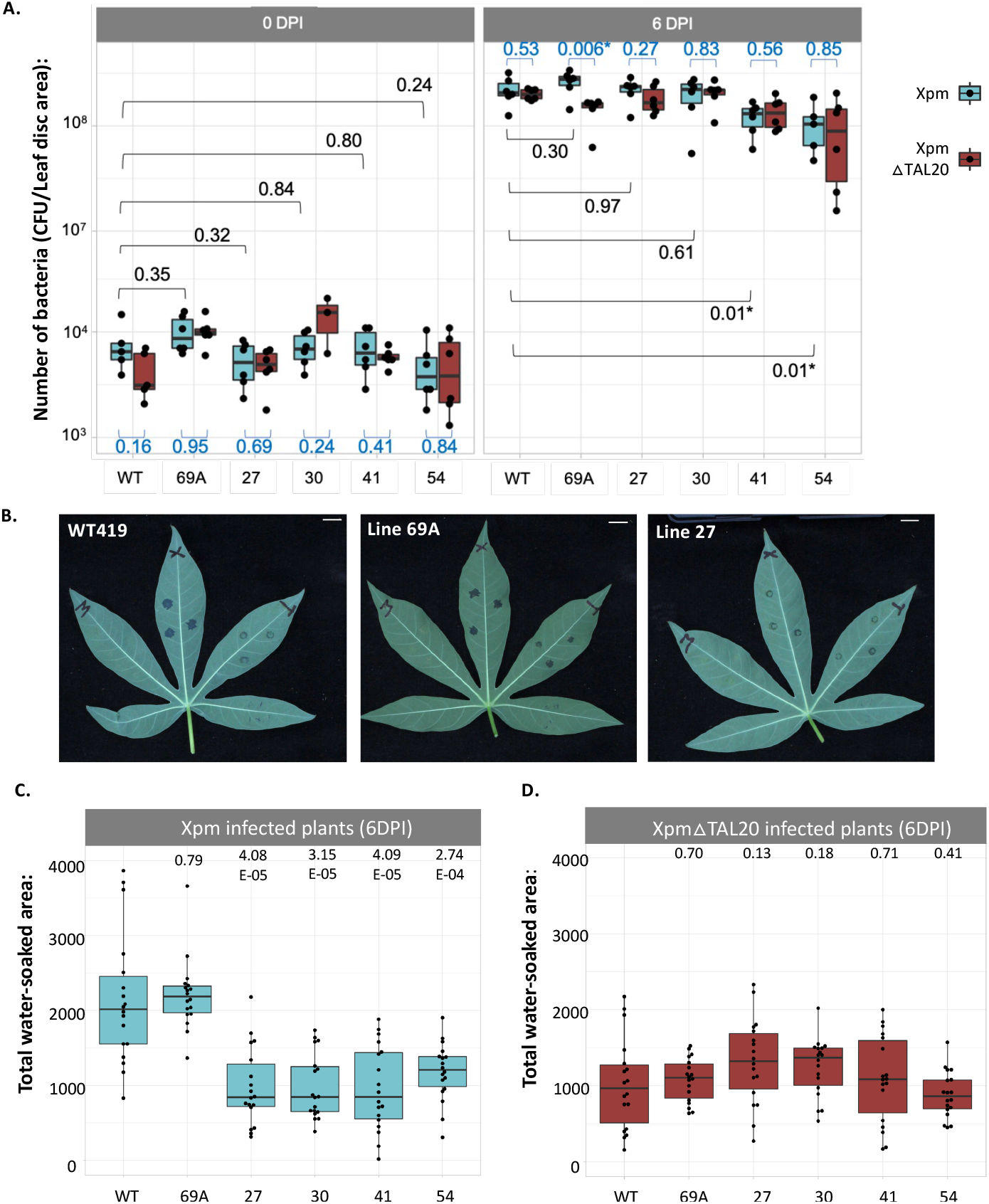
*MeSWEET10a* mutant lines CBB disease symptoms post Xpm infection. **A)** Number of bacteria in cassava leaves measured at 0 days post inoculation (DPI) (left) and 6DPI (right) with Xpm (blue) and Xpm△TAL20 (red) treatments. Colony forming units (CFU/leaf disc area, y-axis) are plotted by plant genotype (x-axis, wildtype or mutant). Black dots represent sample replicates from an independent bacterial growth experiment. **B)** Representative images of infected wildtype (left), silent mutant line #69A (middle), and mutant line #27 (right) cassava leaves detached and imaged at 4DPI. X=Xpm, T=Xpm△TAL20, and M=Mock. Scale bar = 1cm. **C)** Total water-soaked area (pixels, y-axis) of Xpm infected plants (genotypes, x-axis) at 6DPI. **D)** Total water-soaked area (pixels, y-axis) of Xpm△TAL20 infected plants (genotypes, x-axis) at 6DPI. Black dots represent individual water-soaked lesions from three independent water-soaking assay experiments combined. The calculated *p*-values (Unpaired Student’s T-test with unequal variance) are shown above or below each box plot. Black text represents Xpm comparisons between WT and mutant infected plants. Blue text represents comparisons between Xpm and Xpm△TAL20 within each genotype. Dots outside whiskers represent outliers based on default settings of the R package ggplot2. The horizontal line within the box represents the median sample value. The ends of the boxes represent the 3rd (Q3) and 1st (Q1) quartiles. The whiskers show values that are 1.5 times interquartile range (1.5xIQR) above and below Q1 and Q3.

The genotypes of five of these additional lines (**Figure 2B and 2C**) were confirmed using gDNA clone-sequencing to determine mutant line zygosity (**Supplemental Figure 3B**). Line #27 is homozygous with a 185 bp deletion spanning the TATA box, TAL20 binding site, and 5’UTR and a 1 bp frameshift insertion after the start codon. Line #30 is homozygous with a 1 bp deletion upstream of the TATA box and a 2 bp frameshift deletion downstream of the start codon. Line #41 is a biallelic mutant with one allele that has an 11 bp deletion at the TATA box/TAL20 EBE site and a 1 bp insertion in the 5’UTR. The second allele has a 5 bp deletion at the TATA box/TAL20 EBE site and a 1 bp insertion in the 5’UTR. Line #54 is biallelic with one allele containing a 5 bp deletion at the TATA box/TAL20 EBE site and a 1bp insertion in the 5’UTR. The second allele contains 76 bp deletion spanning the TATA box/TAL20 EBE site. Line #69A is a biallelic silent mutant with one allele that has a 1 bp deletion upstream of the TATA box and one insertion in the 5’UTR and another allele that has a 1 bp deletion upstream of the TATA box. gDNA from lines #27, #30, #41, #54, and #69A all produced a *MeSWEET10a* PCR product near 2.1kb corresponding with their insertion/deletion types (**Figure 2D**). An overview of *MeSWEET10a* mutant types generated from all transformations is provided in **Table 1**. Results from select stages of the transformation pipeline are reported in **Supplemental Table 1**. Whole genome sequencing for lines #269, #338, #27, #30, #41, and #54 revealed the transgene insertion number and location for each line (**Supplemental Figure 4**). Mutant plants were moved from tissue culture to soil and phenotypically characterized for height, node number, internode length, petiole length, central lobe length, central lobe width, and whole lobe length (**Figure 2E and Supplemental Table 2:**). Based on these characteristics, mutant plant traits were physiologically similar to wildtype cassava.

**Table 1:** Overview of *MeSWEET10a* mutant line genotypes. Mutation type summary of the 29 transgenic lines recovered from all rounds of transformation based on Sanger sequencing results. ‘*’ denotes a line that died during the tissue culture process.

| Mutant type: | General description: | # of lines: |
| --- | --- | --- |
| WT-like transgenic | Transgenic lines with no edits at gRNA target sites | 1 |
| Silent mutant | Edits that do not impact coding sequence | 13 |
| TAL20 EBE promoter INDEL | Edits within the TAL20 EBE with intact TATA box | 1* |
| *MeSWEET10a* promoter INDEL | Edits in the TATA box predicted to impact expression | 11 |
| *MeSWEET10a* frameshift | Edits after the TSS expected to impact coding sequence | 3 |

**Figure 4:**
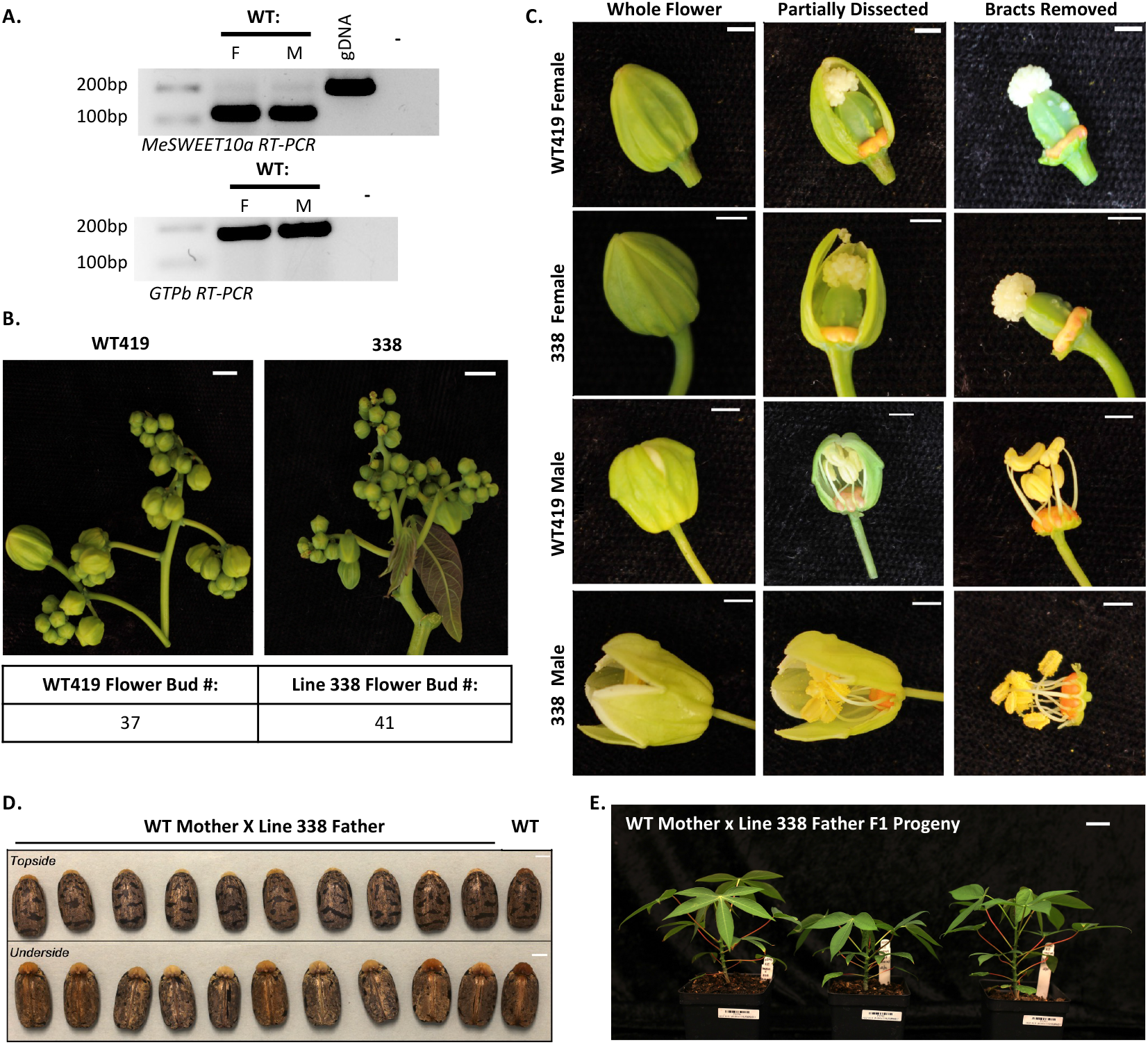
*MeSWEET10a* mutant plant has wildtype-like reproductive morphology and progeny. **A)** RT-PCR of WT419 female (F) and male flowers (M) collected from field grown plants. Top gel shows results of RT-PCR with primers amplifying *MeSWEET10a* with an expected product size of 123 bp. The bottom gel shows results of RT-PCR with primers amplifying the housekeeping gene *GTPb* as a control for sample loading with an expected product size of 184 bp. gDNA is from WT419 leaf tissue included as a positive control and ‘-’ denotes a negative water control. **B)** Representative images of WT419 (left) and line #338 (right) inflorescence structures detached from individual field-grown plants for imaging (top). The number of flower buds present on each inflorescence is presented in table format (bottom). Scale bar = 5cm. **C)**. Representative female and male flowers collected from WT419 and line #338 flowering field-grown plants. Images of the same flower were taken as whole flowers (left) partially dissected with one or two petal-like bracts removed (middle) and dissected with all petal-like bracts removed (right). Scale bar = 0.25cm. **D)** Image of ten seeds derived from WT mother x line #338 father crosses along with an open-pollinated WT419 seed (top). Scale bar = 0.25cm. **E)** Image of three WT419 mother x line #338 father F1 individuals that germinated post-planting in soil.

As a rapid screen to identify mutants that avoid TAL20-mediated induction of *MeSWEET10a*, leaves were detached from plantlets in tissue culture and syringe infiltrated with wildtype Xpm, Xpm△TAL20, or mock treatments. Samples were collected at 48HPI for RNA extraction and RT-PCR analysis (**Supplemental Figure 5**). When infected with Xpm, but not Xpm△TAL20 or mock treatments, the expected 123 bp band corresponding to induction of *MeSWEET10a* was observed in wildtype plants and the silent mutant line #69A. In contrast, no band was observed for samples from lines #27, #30, #41, and #54 (**Figure 2F**). These results support the hypothesis that edits in these lines block induction of *MeSWEET10a* by TAL20.

Xpm and Xpm△TAL20 were infiltrated into cassava leaves and bacterial growth of both strains was similar in infected wildtype and *MeSWEET10a* mutant plants, consistent with our previous research (Veley et al., 2023) (**Figure 3A, Supplemental Figure 6**). However, a visible difference in water-soaking lesions was observed (**Figure 3B, Supplemental infected leaf images available on Figshare)** and these lesions were quantified using a machine learning image analysis method (Elliott et al., 2022). Lines #27, #30, #41, and #54 all exhibited significantly reduced water-soaked lesion area compared to wildtype cassava and line #69A, after challenge with Xpm (**Figure 3C**). There was no significant difference in Xpm△TAL20 lesion area between any of the mutant lines compared to wildtype plants (**Figure 3D**). Similar results were obtained from line #269 and #338 mutant plants infected with Xpm and Xpm△TAL20 (S**upplemental Figure 7**). Therefore, we conclude that *MeSWEET10a* mutant plants have decreased susceptibility to cassava bacterial blight.

Unlike leaves, cassava flowers have endogenous expression of *MeSWEET10a* (Veley et al., 2021). Thus, we wanted to determine if mutating *MeSWEET10a* would impact flower development or reproductive function. Cassava plants do not readily flower and set seed in greenhouse or growth chamber conditions. However, at the time of this study, line #338 had been established in a field site outside Hilo, Hawaii along with WT419 plants. WT419 female and male flower buds were collected for RNA extraction and RT-PCR. Expression of *MeSWEET10a* was confirmed for both flower types (**Figure 4A**). Eleven months after planting, line #338 plants formed the first inflorescences of male and female flower buds (**Figure 4B**). WT419 and line #338 female and male flowers were collected, and the petal-like bracts were dissected (Perera et al., 2012). No obvious visible differences were observed between line #338 and WT419 (**Figure 4C**). Field crosses between WT419 female flowers and line #338 male flowers yielded twelve F1 seeds (**Figure 4D**). Six seeds from WT419 open-pollinated plants were collected for comparison and all seeds were similar in length and width (**Supplemental Figure 8 and Supplemental Table 3**). One common way of measuring seed viability is through float tests, seeds that sink are expected to germinate while seeds that float, commonly do not (Pegman et al., 2017). Three out of twelve WT419 x line #338 seeds sank and as expected, these three seeds successfully germinated, as did two additional seeds that floated. Representative images of F1 plants are provided (**Figure 4E**). The overall germination rate from all WT419 x line #338 F1 seeds was 41.7 percent while the open-pollinated WT419 seeds was 66.7 percent (**Supplemental Table 3**).

## Discussion

This study explored how editing the TAL20 targeted S gene, *MeSWEET10a*, could reduce cassava susceptibility to Xpm-induced bacterial blight. Using a dual gRNA CRISPR/Cas9 strategy, we generated several *MeSWEET10a* mutant lines with edits that impacted the TAL20 EBE binding site and/or the *MeSWEET10a* coding sequence. Mutants that lacked TAL20-mediated induction of *MeSWEET10a* were phenotyped for CBB disease severity through bacterial growth assays and water-soaked lesion analysis. While a consistent difference in bacterial titer was not observed, we found that *MeSWEET10a* mutant lines had significantly reduced water-soaked lesions post Xpm infection compared to infected wildtype cassava. We also inspected the impact of editing *MeSWEET10a* on cassava flowers as the flowers are known to have endogenous expression of the gene (Veley et al., 2021). No obvious defects were observed in *MeSWEET10a* mutants compared to wildtype flowers. Furthermore, field crosses between a *MeSWEET10a* mutant line and WT419 plants produced viable seeds. Overall, these results suggest that editing the *MeSWEET10a* S gene and/or TAL20 EBE site is a viable strategy to improve plant disease resistance and reduce susceptibility to CBB.

The exact mechanics of how *MeSWEET10*a is used by Xpm to promote CBB and pathogen virulence remains unknown. *MeSWEET10*a is a clade III SWEET gene that exports sucrose and glucose from the plant cell into the apoplast where Xpm proliferates. One hypothesis is that Xpm uses these sugars as a carbon source. However, if the *MeSWEET10a* exported sugars were a direct carbon source for Xpm, we would expect that loss of TAL20 would significantly impact bacterial growth. In a previous study, a decrease in bacterial growth was observed for Xpm△TAL20 compared to wildtype Xpm. This previous study used a different genotype of cassava and a modified bacterial growth assay (Cohn and Bart et al., 2014). In the current work, we did not observe a consistently significant difference between Xpm and Xpm△TAL20 colony-forming units (CFUs). Yet, whenever a bacterial growth difference was observed, Xpm△TAL20 titer trended downward compared to Xpm. This may indicate that *MeSWEET10a* has a minor effect on bacterial growth but that the effect is below the limit of sensitivity of the bacterial growth assays and conditions used in our study.

An additional hypothesis is that *MeSWEET10a* exported sugars may serve as an osmolyte for Xpm. As sucrose and glucose are exported out of the plant cell, there is also an osmotic movement of water. *MeSWEET10*a mutants have consistently reduced water-soaked lesions after infection with Xpm compared to infected wildtype cassava. Water-soaked lesions are dark angular spots that occur during pathogenesis as water is moved from the plant cell into the apoplast (Schwartz et al., 2017). Many plant pathogens induce water-soaked leaf lesions during early stages of plant infection (Aung et al., 2018). Other studies have suggested that the role of water-soaking is to create an aqueous environment to aid in bacterial colonization from the plant surface into the apoplast or to help with bacterial spread once *in planta* (Xin et al., 2016). Perhaps the efflux of sugar and water into the apoplast increases bacterial entry into the plant which would not be captured through syringe infiltration-based infection assays. Additionally, Xpm eventually spreads throughout the plant vasculature after initial colonization at the surface of mesophyll cells. It is possible that induction of *MeSWEET10a* by TAL20 may play a role in bacterial spread. Another study in the *Xanthomonas gardneri*-pepper pathosystem reported that reduced water-soaked lesion symptoms did not correlate to a decrease in bacterial growth (Schornack et al., 2008). In the future, additional work is required to tease apart the role of *MeSWEET10a* exported sugars and water-soaking in Xpm pathogenesis.

SWEET genes have been implicated in various roles in plants such as nectar secretion, pollen development, seed filling, and phloem loading (Feng and Frommer, 2015). In cassava, the native function of *MeSWEET10a* remains unknown. However, *MeSWEET10a* is expressed in the cassava flower. The work done with *MeSWEET10a* mutant flowers in this study did not uncover obvious differences between the mutant and wildtype cassava flowers and serves as initial proof of *MeSWEET10a* mutant flower viability. However, further research is needed to determine if editing *MeSWEET10a* has detrimental impacts on flower development and reproductive function. For example, a full field trial with more crosses of the *MeSWEET10a* mutant lines could further validate the ability of mutant plants to produce viable fruit and seed. Additionally, with a larger field trial more seed could be collected to further compare the germination rate of line *MeSWEET10a* mutant plant progeny to WT419 open-pollinated seed. Microscopy comparing the structure of *MeSWEET10a* mutant and wildtype cassava flowers could also determine if there are differences in flower development, not visible to the naked eye.

The ideal *MeSWEET10a* mutant line, for CBB resistance, would contain edits at the TAL20 EBE site while maintaining an intact TATA box and gene coding sequence for native plant function. In this study, one line with EBE only edits and an intact TATA box was recovered, however this line died during the tissue culture process. It is possible that this outcome is selected against for an unknown reason. More likely, if many additional lines were recovered, we would eventually achieve this outcome. One factor complicating the ability to generate EBE specific edits to the *MeSWEET10a* TAL20 binding site is overlap between the TATA box and EBE. In cassava, other TAL effectors localize to EBE sites that include TATA box motifs (Cohn et al., 2016). Other work shows that EBE overlap within or localization near the host TATA box is common in TAL effector S gene target sites [(Grau et al., 2013), (Pereira et al., 2014) and (Pérez-Quintero et al., 2015)]. Alternative gene editing strategies such as base editing or the use of a single CRISPR/Cas9 gRNA for editing at the TAL20 EBE downstream of the TATA box may increase the chances of recovering *MeSWEET10a* mutants with EBE only edits (Azameti and Dauda, 2021). It is also possible that other TAL20 carrying Xpm strains may have effectors with redundant functions and therefore may not exhibit reduced virulence after infection in *MeSWEET10a* mutants. Future work screening *MeSWEET10a* mutants’ susceptibility against other Xpm isolates would help determine the overall viability of a disease resistance strategy based on editing the *MeSWEET10a* S gene alone. Likewise, developing *MeSWEET10a* mutants in additional cassava cultivars and examining CBB susceptibility would be useful. It is important to note that editing one S gene may not be enough to significantly reduce cassava susceptibility to CBB in a field setting. Thus, investigating the role of additional S genes on Xpm virulence and developing cassava mutants with stacked edits at different S gene targets may be required to develop plants with sustained resistance to CBB.

## Materials and Methods

### Construct design and cloning

The *MeSWEET10a* (Manes.06G123400) FASTA sequence file was downloaded from Phytozome (*Manihot esculenta* genome v6.1) and uploaded to the software Geneious. Notable promoter regulatory elements were annotated as previously reported by Cohn and Bart et. al., 2014 including the effector binding element (EBE) site where TAL20 binds. The reported EBE sequence was confirmed using the TALEnt target finder tool. The Geneious “find CRISPR sites” function was used to find all potential targets and Cas9 *(S. pyogenes*)-specific PAM sites (sequence: 5′-NGG-3′). Candidate guide RNA (gRNAs) target sequences were selected based on those whose targets were near the TAL20 EBE and the translational start site of *MeSWEET10a* or within the 5’UTR. Candidate gRNAs were further analyzed by comparing the gRNAs against the cassava genome to identify potential off-targets using NCBI-BLAST. All constructs were assembled using a multiple gRNA spacer Csy4 array as previously described (Čermák et al., 2017). Three constructs were used for this study. All constructs were designed to carry two gRNAs, were cloned in the pTRANS_220D backbone, and have a kanamycin resistance cassette. Construct 108 carries gRNA1 (GAGAAGCGTTTATATAGGGG) which targets the TAL20 EBE site and gRNA2 (GAAGTCCAATGACAAGTGCA) targeting the *MeSWEET10a* translation start site. Construct 108 also carries an intended EBE repair template (as an attempt to replace the sequence) containing homology arms that flank the EBE (1,079bp 5’ homology arm and 727 bp 3’ homology arm). Construct 249 was designed to carry the TAL20 EBE site target gRNA1 (GAGAAGCGTTTATATAGGGG) and gRNA3 (ACTCTCTTTCCCTTGTGCAG) which targets the 5’UTR with no repair template. Construct 250 was designed to carry gRNA4 (AAAATATGTCAATGTAACAG) and gRNA5 (TATGTTGTGCAATGATGGAT) which target the 5’UTR and EBE with no repair template. Construct assembly was confirmed through colony PCR, Sanger sequencing, and by Illumina sequencing. Constructs were transformed into LBA4404 Agrobacterium cells for cassava transformations. All construct sequences, maps, and Illumina reads are available in supplementary data.

### Plant materials and growing conditions

Transgenic cassava lines expressing the CRISPR/Cas9 machinery and gRNAs were generated in the cassava cultivar TME419 (WT419) through *Agrobacterium*-mediated transformation as described (Chauhan et al., 2015). Transgenic FEC cells were selected for resistance using 100mM paramomycin (275uL/L) on spread plates. 100mM paramomycin (450uL/L) was used to further select for resistant transgenic cells on stage 1, 2, and 3 plates. Transgenic FEC cells and eventual transgenic plantlets were maintained in tissue culture in conditions set to 28°C +/-1°C, 75 μmol·m^-2^ ·s^-1^ light; 16 hrs light / 8 hrs dark. Plantlets were transferred to soil on a misting bench and covered with domes to maintain high humidity. After establishment in soil, plants were moved from the misting bench and acclimated to greenhouse conditions set to 28°C; 50% humidity; 16 hrs light / 8 hrs dark, and 1000 W light fixtures that supplemented natural light levels below 400 W / m^2^. Following bacterial infection assays, plants were kept in a post-treatment growth chamber with conditions set to 27°C; 50% humidity, and 12 hrs light / 12 hrs dark. F1 seeds generated from WT419 × 338 crosses were planted in soil and kept in a plant growth chamber set to 37°C, 60% humidity, 12 hrs light / 12 hrs dark at 400 μmol·m^-2^·s^-1^. Once seedlings germinated, they were transferred to larger pots and moved to greenhouse conditions listed above.

### DNA extraction and transgenic line genotyping

As an initial pass: transgenic lines recovered from the first transformation with construct 108, a PCR followed by restriction digest and gel electrophoresis strategy was used. Mutants with varying HaeIII digest patterns were suspected to have deletions and were moved forward for Sanger sequencing. In subsequent rounds of transformation, mutants with both point mutations and insertions/deletions were further characterized.

For later transformations: leaf lobe samples from transgenic lines were collected from 2-3 individual plantlets and pooled into 2mL Eppendorf Safelock tubes with three disposable 3 mm Propper solid glass beads. The sample tubes were flash frozen in liquid nitrogen and ground to a fine powder using a QIAGEN TissueLyser II machine at 30hz for 3 minutes until the sample was fully homogenized. Genomic DNA was extracted using the Sigma GenElute Plant Genomic DNA Miniprep Kit. The *MeSWEET10a* region of interest was amplified using “outer” primers designed to avoid amplification of the EBE repair template present in the construct 108 transgene and a 2.1kb product was generated for each line. All primers used in this study are provided in **Supplemental Table 4**. The PCR product was purified using the Qiagen QIAquick PCR Purification kit. The samples were sent for Sanger sequencing using secondary “inner” primers designed to start amplification closer to the gRNA target sites. Transgenic line trace files were compared to wildtype TME419 trace files and edits within and across each gRNA were identified using the Geneious bioinformatics tool. Outer and inner primer sequences used for genotyping are available in the primer list table. Clone-seq was performed on select lines to determine if edits were homozygous for each allele.

### Identification of transgene location(s)

For each construct a custom reference genome was created which contained the haplotype-resolved genome assembly for cassava variety TME204 (Qi et al., 2022) along with the vector sequence from the T-DNA left border through the T-DNA right border of the appropriate construct. The program bwa mem (version 0.7.12-r1039) was used to align the whole genome sequencing data to the custom reference genome (Li, 2013). Illumina reads where one pair aligned to the T-DNA insertion sequence and the other aligned to the cassava genome were isolated using samtools (version 1.11) (Danecek et al., 2021) and used as input for *de novo* assembly by Trinity (version v2.1.1) (Grabherr et al., 2011). Resultant contigs were then used in a blastn (version 2.12.0+) query against a BLAST database of the custom reference genome initially used for bwa alignments (Sayers et al., 2022). Contigs where a portion matched the cassava genome and another portion matched the T-DNA insertion sequence identified the coordinates of the 5’ and 3’ ends of an insertion point within the genome. Integrative Genomics Viewer (IGV; version 2.12.3) was used for manual inspection and visualization of the aligned WGS data to the custom T-DNA insertion plus genome (Thorvaldsdóttir et al., 2013).

### Bacterial inoculations

*Xanthomonas* strains were struck from glycerol stocks onto NYG agar plates containing appropriate antibiotics. The strains used were Xpm668 (rifampicin 50 μg/ml), Xpm668ΔTAL20 (suicide vector knockout, tetracycline 5 μg/ml, rifampicin 50 μg/ml), Xe85-10 (rifampicin 50 μg/ml) and Xe85-10+TAL20_Xpm668_ (rifampicin 50 μg/ml, kanamycin 50 μg/ml), (Cohn and Bart et al., 2014). *Xanthomonas* strains were grown in a 30°C incubator for 2-3 days. Inoculum for each strain was made by transferring bacteria from plates into 10mM MgCl_2_ using inoculation loops and brought up to a concentration of OD600 = 0.01 for bacterial growth and water-soaked lesion assays and OD600 = 1 for RT-PCR. Leaves on cassava plants were inoculated using a 1.0 mL needleless syringe. For each replicate assay, two cassava plants per background (WT or transgenic) were used for inoculations, and four leaves were inoculated on each plant. One bacterial strain suspended in 10mM MgCl_2_ was inoculated per leaf lobe with three injection sites and mock inoculations of 10mM MgCl_2_ alone were included. In total, there were nine infiltrated sites per leaf.

### RNA extraction and RT-PCR analysis

For lines with edits of interest, RNA extraction and RT-PCR was performed at the plantlet stage in tissue culture and on soil established plants in the greenhouse. At the plantlet stage, 9 leaves were detached from every transgenic line and 3 leaves each were syringe infiltrated with either mock (10 mM MgCl_2_alone) or *Xanthomonas* (*Xpm668*) on sterile petri dishes. For each line, a set of 3 infiltrated leaves per treatment were kept on MS2 plates in a post-treatment room light shelf. At 48 hours post infection, samples were collected, and 3 infiltrated leaves per treatment were pooled into Eppendorf safelock tubes with 3mm glass beads. For greenhouse plants, one leaf was selected (3 biological replicates per plant background) and syringe infected with 3 infiltrated sites per treatment (mock (10 mM MgCl_2_ alone) or *Xanthomonas* (*Xpm668*) +/-TAL20 strains, or *Xanthomonas euvesicatoria* (Xe85-10) +/-TAL20 strains) on separate leaf lobes. At 48HPI, leaf punches from the infiltrated sites were collected using a size 7 mm core borer and technical reps were pooled into Eppendorf safelock tubes with 3mm glass beads. Samples were flash frozen in liquid nitrogen and ground using TissueLyser settings described above. Total RNA was extracted from each sample using the Sigma Spectrum Plant Total RNA Kit. 1 μg of RNA was DNase treated using Promega RQ1 DNase enzyme and reverse transcribed into cDNA using Thermo Fisher Scientific SuperScript III Reverse Transcriptase. RT-PCR was performed on each sample using primers specific to *MeSWEET10a* and to cassava *GTPb* (Manes.09G086600) as a constitutively expressed control. All primers used in this study are provided in **Supplemental Table 4**. RT-PCR results were analyzed to identify transgenic lines in which ectopic expression of *MeSWEET10a* was not induced by *Xanthomonas* (+TAL20) infection as is normally seen in wildtype cassava infected with *Xanthomonas* (+TAL20).

### Bacterial growth assay

Cassava leaves were infiltrated with either 10 mM MgCl_2_ (mock control) or *Xanthomonas* (*Xpm668* strains +/-TAL20) suspended in 10 mM MgCl_2_ as described above. Leaf punch samples were taken at the site of infiltration using a 7 mm cork borer (size 4) at 0-, 2-, 4-and 6-days post inoculation. For day-0 samples, infiltrated spots were allowed to dry down prior to processing. Individual leaf punches were transferred to 2 mL Eppendorf Safelock tubes with 200 uL of 10 mM MgCl_2_ and three disposable 3 mm Propper solid glass beads. Samples were ground with a Qiagen Tissuelyzer at 28 hZ for 3 minutes. 200 uL of the ground sample was transferred to the first column of a labeled 96-well plate. Serial dilutions were performed by transferring 20 uL of the non-dilute sample (10^1^) to the next well containing 180 ul of 10 mM MgCl_2_. Samples were serially diluted to 10^4^for day 0, 10^6^ for day 2 and 4, and for 10^8^ days 6. 10 ul of each serial dilution was pipetted and spread onto labeled quadrants of an NYG plate with cycloheximide and the appropriate antibiotics for the infiltrated bacterial strain. Plates were incubated at 30°C for 2-3 days and the number of colonies were counted. Colony Forming Units (CFU) reported in this manuscript were transformed by sample area (CFU/leaf disc area where disc area = 0.38cm).

### Water-soaked lesion imaging and quantification

Cassava leaves were detached and imaged at 0-, 6-, and 9-days post inoculation (DPI). One leaf for every plant was collected for a total of two leaves per plant background at each time point. Line 338 and 269 leaves were imaged from above using a Raspberry Pi Sony IMX219 camera in an enclosed box with an overhead light. To increase image resolution, all subsequent infected plant leaves were imaged from above using a Canon EOS Rebel T5i camera with a 15-85mm lens in an enclosed box with an overhead light. Images were processed and analyzed for water-soaked lesion area and gray-scale color using a previously described custom machine learning image analysis tool developed for CBB disease quantification (Elliott et al., 2022).

### Flower inflorescences and flower bud imaging and dissection

Flower inflorescences and individual buds were detached from cassava plants (WT419 or line 338) growing in a field site (Hilo, Hawaii, USA). All flower inflorescences and individual buds were imaged in the field from above using a Canon EOS Rebel T5i camera with a 15-85mm lens in a portable, partially enclosed pop-up light box with built in LED lights controlled by a USB power pack. Images were post processed using photoshop for color correction and a scale bar was added using ImageJ version FIJI.

## Author Contributions

KE, KMV, and RSB designed the study. KMV designed the constructs. GJ and KMV generated the first generation *MeSWEET10a* mutant (lines #338 and #269) and completed RT-PCR analysis on them. KE completed transformations and genotyping for all additional *MeSWEET10a* mutant lines. KE completed bacterial infections, RT-PCR, bacterial growth assays, and water-soaked lesion analyses. KBG completed transgene insertion analysis on *MeSWEET10a* mutant lines. AW collected plant trait measurements for *MeSWEET10a* mutant and WT419 plants. JN and LK managed the cassava field and performed all crosses. KE collected, dissected, and imaged all male and female flowers. KE and MY collected line F1 seed measurements and tested the germination rate. KE wrote the original manuscript draft, completed statistical analysis, and generated figures. KE, KMV, and RSB assisted in manuscript and figure review and editing. RSB provided supervision over the project and SMW provided supervision over field work. All authors read and approved the final manuscript.

## Acknowledgments

We would like to acknowledge and thank the current and former Bart Lab members who provided insightful discussion and feedback on this project, especially Dr. Daniel Lin, Dr. Qi Wang, Taylor Harris, Jeffrey C. Berry, and Joshua Sumner. Thank you to Dr. Nigel Taylor and members of his lab who provided the Friable Embryogenic Callus used for cassava transformation and transformation expertise.

## Data Availability

The authors confirm that experimental data are available and accessible via the main text and/or the supplemental data. All raw data, R scripts, and water-soaked lesion images are available on figshare: 10.6084/m9.figshare.22718680

## Supplemental Figure Legends

**Supplemental Figure 1**: Construct 108 transgenic lines restriction digest screen **A)** Overview of HaeIII restriction digest strategy for *MeSWEET10a* region of interest in WT (top) and potential mutant with the EBE template repair (bottom). WT-like sequences were expected to remain uncut. However, mutants with integration of the repair template were expected to have an abolished HaeIII site and remain uncut. Blue arrows point to the HaeIII cut site and potential gRNA2 repair site from the template integration by homology-directed repair. **B)** Example of an HaeIII digest performed on sixteen transgenic lines. White arrows point to digest patterns unlike the WT digest pattern. The expected digest pattern for WT is bands at 895, 384, 270, 271, 195, and 192 bp. Bolded line numbers (#2, #269, and #338) were moved forward for Sanger sequencing.

**Supplemental Figure 2**: Transgenic line sequencing from first construct 108 transformation **A)** Geneious screenshots showing Sanger-sequencing results for lines #2, #243, and #323. Mutation types and INDELs are described in text. **B)** Geneious screenshots of Sanger-sequencing results of *E. coli* clones containing *MeSWEET10a* gDNA from lines #269 and #338 plants.

**Supplemental Figure 3**: Constructs 108, 249, and 250 Sanger-sequencing data from four replicate transformations **A)** Geneious screenshots showing Sanger-sequencing results for all lines obtained by additional transformation with constructs 108, 249, and 250. Mutation types and INDELs are described in text. ‘*’ Denotes a line with EBE specific mutants that did not survive in tissue culture. **B)** Geneious screenshots of Sanger-sequencing results of *E. coli* clones containing *MeSWEET10a* gDNA from lines #27, #30, #41, #54, and #69A plants.

**Supplemental Table 1**: Transformation results overview Table with summary of transformation recovery statistics from all rounds of transformation separated by construct type. Callus selected represents the number of lines that survived selection and made it to stage 2 plates for the cassava transformation pipeline. Mature lines recovered represent transgenic lines that survived as plantlets up to the first MS2 plate stage of cassava transformation as described by Chauhan et al (Chauhan et al., 2015). Select lines were characterized and the number of lines with edits are reported.

**Supplemental Table 2**: Plant morphology measurements Measurements of cassava morphology traits including plant height (cm), node numbers, internode lengths above and below the woody transition (cm), leaf lobe number, central lobe length and width(cm), and whole length width (cm) for 3 representative individuals each of WT419 and lines #27, #30, #41, and #54 plants. Line #69A had 2 individual plants. The average of each trait measurement was taken along with standard deviation (measurement ± standard deviation). The calculated p-value from unpaired student T-tests with unequal variance is shown for each measurement comparing WT and each mutant line.

**Supplemental Figure 4**: Transgene insertion analysis Graphic depicting transgene insertion number and location in lines #269, #338, #27, #30, #41, and #54. For each line, the total number of insertions is listed (left-hand side). For each insertion, the haplotype and chromosome location for the site of insertion is noted. Red triangles and brackets depict the site of transgene insertion. The bracket indicates a larger distance between insertion coordinates. Nearby genes are annotated with their Manes ID.

**Supplemental Figure 5**: Xpm infected detached leaf RT-PCR RT-PCR of wildtype (WT419) cassava and *MeSWEET10a* mutant lines (#69A, #69B, #1-18, #11, #18, #21, #27, #30, #35, #41, #49, #54, #88 and #108) detached leaves from plantlets infected with Xpm. The top gel shows results of RT-PCR with primers amplifying *MeSWEET10a* with an expected product size of 123 bp. The bottom gel shows results of RT-PCR with primers amplifying the housekeeping gene *GTPB* as a control for sample loading with an expected product size of 184 bp. ‘+’ denotes a WT419 gDNA positive control. ‘-’ denotes a negative water control. The bold text represents mutants that lack TAL20 mediated induction of *MeSWEET10a*.

**Supplemental Figure 6**: Additional replicates of bacterial growth assays **A)** Replicate 1 bacterial growth assay for WT, line #69A, #30, #41, and #54 plants. Number of bacteria in cassava leaves measured at 0, 2, 4, and 6DPI post syringe infiltration with Xpm (blue) and Xpm△TAL20 (red) treatments. **B)** Replicate 1 bacterial growth assay for mutant line 27. Along with WT and #69A controls. Number of bacteria in cassava leaves measured at 0, 2, 4, and 6DPI post syringe infiltration with Xpm (blue) and Xpm△TAL20 (red) treatments. **C)** Replicate 2 bacterial growth assay for WT, line #69A, #27, #30, #41, and #54 plants. Number of bacteria in cassava leaves measured at 0 (left) and 6DPI (right) post syringe infiltration with Xpm (blue) and Xpm△TAL20 (red) treatments. For all box plots, Colony Forming Units (CFU/cm2, Y-axis) are plotted by plant genotype (X-axis) tested (wildtype or mutant). Black dots represent technical replicates from one independent bacterial growth experiment. Results of statistical analyses (Unpaired student’s t-test with unequal variance) comparing the difference between Xpm growth across wild-type and mutant cassava genotypes infected with Xpm. P-values are shown above or below brackets indicating the comparison types for statistical analyses. Black dots represent individual water-soaked lesions from three independent water-soaking assay experiments combined. In all boxplots, the calculated *p*-values (Unpaired Student’s T-test with unequal variance) are shown above or below each box plot. Black text represents Xpm comparisons between WT and mutant infected plants. Blue text represents comparisons between Xpm and Xpm△TAL20 within each genotype. Dots outside whiskers represent outliers based on default settings of the R package ggplot2. The horizontal line within the box represents the median sample value. The ends of the boxes represent the 3rd (Q3) and 1st (Q1) quartiles. The whiskers show values that are 1.5 times interquartile range (1.5xIQR) above and below Q1 and Q3.

**Supplemental Figure 7**: Lines #269 and #338 water-soaked lesion assay **A)** Representative images of infected wildtype (left) and mutant line #338 (right) cassava leaves detached from the plant and imaged at 6DPI. X= Xpm, T=Xpm△TAL20, and M=Mock. Scale bar = 1cm. **B)** Total water-soaked area (pixels, y-axis) of mock and Xpm infected plants (genotypes, x-axis) at 9DPI. **C)** Total water-soaked area (pixels, y-axis) of mock and Xpm△TAL20 infected plants (genotypes, x-axis) at 9DPI. Black dots represent individual water-soaked lesions from three independent water-soaking assay experiments combined. Calculated *p*-values (Unpaired Student’s T-test with unequal variance) comparing mutant line to wildtype water-soaked area shown above each box plot. For all box plots, dots outside whiskers represent outliers. The horizontal line within the box represents the median sample value. The ends of the boxes represent the 3rd (Q3) and 1st (Q1) quartiles. The whiskers show values that are 1.5 times interquartile range (1.5xIQR) above and below Q1 and Q3.

**Supplemental Figure 8**: WT419 x #338 F1 Seed Traits The measurements recorded from twelve F1 seeds (A-L) recovered from crosses of WT419 mother by #338 father flowers. Measurements include **A)** Seed length (cm) **B)** seed width (cm), and **C)** seed weight (mg).

**Supplemental Table 3**: Recovered seed information Table of recorded information for WT419 by #338 F1 seed and WT419 open pollinated seed recovered from the field and tested for germination. Seed ID, Letter ID, Date of Cross, Mother, Father, Seed length (cm), width (cm), weight (mg), Float test results, and germination data are reported. Results for seed float test denoted as ‘s’ for sink and ‘f’ for float.

**Supplemental Table 4**: Primer list Table of all primers used in this study. The primer name/stock number (left), sequence (middle), and description (right) are provided.

## References

Abrusci P, McDowell MA, Lea SM, Johnson S (2014) Building a secreting nanomachine: a structural overview of the T3SS. Current Opinion in Structural Biology 25: 111–117

Alves AAC (2002) Cassava botany and physiology. In RJ Hillocks, JM Thresh, eds, Cassava: biology, production and utilization. CABI, Wallingford, pp 67–89

An S-Q, Potnis N, Dow M, Vorhölter F-J, He Y-Q, Becker A, Teper D, Li Y, Wang N, Bleris L, et al Mechanistic insights into host adaptation, virulence and epidemiology of the phytopathogen Xanthomonas. FEMS Microbiol Rev. doi: 10.1093/femsre/fuz024

Antony G, Zhou J, Huang S, Li T, Liu B, White F, Yang B (2010) Rice xa13 Recessive Resistance to Bacterial Blight Is Defeated by Induction of the Disease Susceptibility Gene Os-11N3. The Plant Cell 22: 3864–3876

Aung K, Jiang Y, He SY (2018) The role of water in plant–microbe interactions. The Plant Journal 93: 771–780

Azameti MK, Dauda WP (2021) Base Editing in Plants: Applications, Challenges, and Future Prospects. Frontiers in Plant Science 12:

Bart R, Cohn M, Kassen A, McCallum EJ, Shybut M, Petriello A, Krasileva K, Dahlbeck D, Medina C, Alicai T, et al (2012) High-throughput genomic sequencing of cassava bacterial blight strains identifies conserved effectors to target for durable resistance. Proc Natl Acad Sci USA 109: E1972–1979

Boch J, Bonas U (2010) Xanthomonas AvrBs3 family-type III effectors: discovery and function. Annu Rev Phytopathol 48: 419–436

Britt AB, May GD (2003) Re-engineering plant gene targeting. Trends Plant Sci 8: 90–95

Cernadas RA, Doyle EL, N. ño-Liu DO, Wilkins KE, Bancroft T, Wang L, Schmidt CL, Caldo R, Yang B, White FF, et al (2014) Code-assisted discovery of TAL effector targets in bacterial leaf streak of rice reveals contrast with bacterial blight and a novel susceptibility gene. PLoS Pathog 10: e1003972

Chauhan RD, Beyene G, Kalyaeva M, Fauquet CM, Taylor N (2015) Improvements in Agrobacterium-mediated transformation of cassava (Manihot esculenta Crantz) for large-scale production of transgenic plants. Plant Cell, Tissue and Organ Culture (PCTOC) 121: 591–603

Chen L-Q (2014) SWEET sugar transporters for phloem transport and pathogen nutrition. New Phytologist 201: 1150–1155

Chen L-Q, Hou B-H, Lalonde S, Takanaga H, Hartung ML, Qu X-Q, Guo W-J, Kim J-G, Underwood W, Chaudhuri B, et al (2010) Sugar transporters for intercellular exchange and nutrition of pathogens. Nature 468: 527–532

Cohn M, Bart RS, Shybut M, Dahlbeck D, Gomez M, Morbitzer R, Hou B-H, Frommer WB, Lahaye T, Staskawicz BJ (2014) Xanthomonas axonopodis virulence is promoted by a transcription activator-like effector-mediated induction of a SWEET sugar transporter in cassava. Mol Plant Microbe Interact 27: 1186–1198

Cohn M, Morbitzer R, Lahaye T, Staskawicz BJ (2016) Comparison of gene activation by two TAL effectors from Xanthomonas axonopodis pv. manihotis reveals candidate host susceptibility genes in cassava. Molecular Plant Pathology 17: 875–889

Constantin EC, Cleenwerck I, Maes M, Baeyen S, Van Malderghem C, De Vos P, Cottyn B (2016) Genetic characterization of strains named as Xanthomonas axonopodis pv. dieffenbachiae leads to a taxonomic revision of the X. axonopodis species complex. Plant Pathology 65: 792–806

Cox KL, Meng F, Wilkins KE, Li F, Wang P, Booher NJ, Carpenter SCD, Chen L-Q, Zheng H, Gao X, et al (2017) TAL effector driven induction of a SWEET gene confers susceptibility to bacterial blight of cotton. Nature Communications 8: 1–14

Danecek P, Bonfield JK, Liddle J, Marshall J, Ohan V, Pollard MO, Whitwham A, Keane T, McCarthy SA, Davies RM, et al (2021) Twelve years of SAMtools and BCFtools. GigaScience 10: giab008

Eckardt NA (2002) Plant Disease Susceptibility Genes? The Plant Cell 14: 1983–1986

Elliott K, Berry JC, Kim H, Bart RS (2022) A comparison of ImageJ and machine learning based image analysis methods to measure cassava bacterial blight disease severity. Plant Methods 18: 86

EL-Sharkawy MA (2003) Cassava biology and physiology. Plant Mol Biol 53: 621–641

Feng L, Frommer WB (2015) Structure and function of SemiSWEET and SWEET sugar transporters. Trends Biochem Sci 40: 480–486

Grabherr MG, Haas BJ, Yassour M, Levin JZ, Thompson DA, Amit I, Adiconis X, Fan L, Raychowdhury R, Zeng Q, et al (2011) Full-length transcriptome assembly from RNA-Seq data without a reference genome. Nat Biotechnol 29: 644–652

Grau J, Wolf A, Reschke M, Bonas U, Posch S, Boch J (2013) Computational Predictions Provide Insights into the Biology of TAL Effector Target Sites. PLOS Computational Biology 9: e1002962

Gupta PK, Balyan HS, Gautam T (2021) SWEET genes and TAL effectors for disease resistance in plants: Present status and future prospects. Mol Plant Pathol 22: 1014–1026

Hillocks RJ, Thresh JM, Bellotti A (2002) Cassava: Biology, Production and Utilization. CABI

Hogenhout SA, Van der Hoorn RAL, Terauchi R, Kamoun S (2009) Emerging Concepts in Effector Biology of Plant-Associated Organisms. MPMI 22: 115–122

Hu Y, Zhang J, Jia H, Sosso D, Li T, Frommer WB, Yang B, White FF, Wang N, Jones JB (2014) Lateral organ boundaries 1 is a disease susceptibility gene for citrus bacterial canker disease. Proc Natl Acad Sci USA 111: E521–529

Hu Z, Tang Z, Yang J, Bao S, Zhang Y, Ma L, Zheng Q, Yang F, Zhang D, Sun S, et al (2023) Knockout of OsSWEET15 Impairs Rice Embryo Formation and Seed-Setting. Plant and Cell Physiology 64: 258–268

Huang X, Wang Y, Wang N (2022) Highly Efficient Generation of Canker-Resistant Sweet Orange Enabled by an Improved CRISPR/Cas9 System. Frontiers in Plant Science 12:

Jacques M-A, Arlat M, Boulanger A, Boureau T, Carrère S, Cesbron S, Chen NWG, Cociancich S, Darrasse A, Denancé N, et al (2016) Using Ecology, Physiology, and Genomics to Understand Host Specificity in Xanthomonas. Annual Review of Phytopathology 54: 163–187

Kandel SL, Joubert PM, Doty SL (2017) Bacterial Endophyte Colonization and Distribution within Plants. Microorganisms 5: 77

Leyns F, Cleene MD, Swings J-G, De Ley J (1984) The Host Range of the Genus Xanthomonas. Botanical Review 50: 308–356

Li H (2013) Aligning sequence reads, clone sequences and assembly contigs with BWA-MEM. doi: 10.48550/arXiv.1303.3997

López CE, Bernal AJ (2012) Cassava bacterial blight: using genomics for the elucidation and management of an old problem. Tropical Plant Biology 5: 117–126

Lozano JC (1986) Cassava Bacterial Blight: A Manageable Disease. Plant Dis 70: 1089

Lozano JC, Byrne D, Bellotti A (1980) Cassava/Ecosystem Relationships and their Influence on Breeding Strategy. Tropical Pest Management 26: 180–187

Mhedbi-Hajri N, Hajri A, Boureau T, Darrasse A, Durand K, Brin C, Saux MF-L, Manceau C, Poussier S, Pruvost O, et al (2013) Evolutionary History of the Plant Pathogenic Bacterium Xanthomonas axonopodis. PLOS ONE 8: e58474

Morgan NK, Choct M (2016) Cassava: Nutrient composition and nutritive value in poultry diets. Animal Nutrition 2: 253–261

Moscou MJ, Bogdanove AJ (2009) A simple cipher governs DNA recognition by TAL effectors. Science 326: 1501

Oliva R, Ji C, Atienza-Grande G, Huguet-Tapia JC, Perez-Quintero A, Li T, Eom J-S, Li C, Nguyen H, Liu B, et al (2019) Broad-spectrum resistance to bacterial blight in rice using genome editing. Nat Biotechnol 37: 1344–1350

Pegman APM, Perry GLW, Clout MN (2017) Size-based fruit selection by a keystone avian frugivore and effects on seed viability. New Zealand Journal of Botany 55: 118–133

Pereira AL, Carazzolle MF, Abe VY, de Oliveira ML, Domingues MN, Silva JC, Cernadas RA, Benedetti CE (2014) Identification of putative TAL effector targets of the citrus canker pathogens shows functional convergence underlying disease development and defense response. BMC Genomics 15: 157

Perera PIP, Quintero M, Dedicova B, Kularatne JDJS, Ceballos H (2012) Comparative morphology, biology and histology of reproductive development in three lines of Manihot esculenta Crantz (Euphorbiaceae: Crotonoideae). AoB Plants 5: pls046

Pérez-Quintero A, Lamy L, Gordon J, Escalon A, Cunnac S, Szurek B, Gagnevin L (2015) QueTAL: a suite of tools to classify and compare TAL effectors functionally and phylogenetically. Frontiers in Plant Science 6:

Phillips AZ, Berry JC, Wilson MC, Vijayaraghavan A, Burke J, Bunn JI, Allen TW, Wheeler T, Bart RS (2017) Genomics-enabled analysis of the emergent disease cotton bacterial blight. PLOS Genetics 13: e1007003

Puchta H (2005) The repair of double-strand breaks in plants: mechanisms and consequences for genome evolution. Journal of Experimental Botany 56: 1–14

Qi W, Lim Y-W, Patrignani A, Schläpfer P, Bratus-Neuenschwander A, Grüter S, Chanez C, Rodde N, Prat E, Vautrin S, et al (2022) The haplotype-resolved chromosome pairs of a heterozygous diploid African cassava cultivar reveal novel pan-genome and allele-specific transcriptome features. Gigascience 11: giac028

Ryan RP, Vorhölter F-J, Potnis N, Jones JB, Van Sluys M-A, Bogdanove AJ, Dow JM (2011) Pathogenomics of Xanthomonas: understanding bacterium–plant interactions. Nature Reviews Microbiology 9: 344–355

Sayers EW, Bolton EE, Brister JR, Canese K, Chan J, Comeau DC, Connor R, Funk K, Kelly C, Kim S, et al (2022) Database resources of the national center for biotechnology information. Nucleic Acids Research 50: D20–D26

van Schie CCN, Takken FLW (2014) Susceptibility Genes 101: How to Be a Good Host. Annual Review of Phytopathology 52: 551–581

Schornack S, Minsavage GV, Stall RE, Jones JB, Lahaye T (2008) Characterization of AvrHah1, a novel AvrBs3-like effector from Xanthomonas gardneri with virulence and avirulence activity. New Phytologist 179: 546–556

Schornack S, Moscou MJ, Ward ER, Horvath DM (2013) Engineering Plant Disease Resistance Based on TAL Effectors. Annual Review of Phytopathology 51: 383–406

Schwartz AR, Morbitzer R, Lahaye T, Staskawicz BJ (2017) TALE-induced bHLH transcription factors that activate a pectate lyase contribute to water soaking in bacterial spot of tomato. Proceedings of the National Academy of Sciences 114: E897–E903

Taylor N, Chavarriaga P, Raemakers K, Siritunga D (2003) Development and application of transgenic technologies in cassava.

Thorvaldsdóttir H, Robinson JT, Mesirov JP (2013) Integrative Genomics Viewer (IGV): high-performance genomics data visualization and exploration. Briefings in Bioinformatics 14: 178–192

Veley KM, Elliott K, Jensen G, Zhong Z, Feng S, Yoder M, Gilbert KB, Berry JC, Lin Z-JD, Ghoshal B, et al (2023) Improving cassava bacterial blight resistance by editing the epigenome. Nat Commun 14: 85

Veley KM, Okwuonu I, Jensen G, Yoder M, Taylor NJ, Meyers BC, Bart RS (2021) Gene tagging via CRISPR-mediated homology-directed repair in cassava. G3 (Bethesda) 11: jkab028

Xin X-F, Nomura K, Aung K, Velásquez AC, Yao J, Boutrot F, Chang JH, Zipfel C, He SY (2016) Bacteria establish an aqueous living space in plants crucial for virulence. Nature 539: 524–529

